# Evidence for gene transfer between mycoviruses and their host: *Curvulaviridae* as a case study

**DOI:** 10.1101/2023.07.20.549826

**Authors:** Ayoub Maachi, Pau Alfonso, Esmeralda G. Legarda, Beilei Wu, Santiago F. Elena

**Affiliations:** Instituto de Biología Integrativa de Sistemas (CSIC-Universitat de València), Paterna, 46980 Valencia, Spain; Institute of Plant Protection, Chinese Academy of Agricultural Sciences, Haidian, 100193 Beijing, China; Santa Fe Institute, Sant Fe, NM 87501, USA

**Keywords:** coat protein, fungal viruses, molecular evolution, origin of viral genes, phylogenetics, virus evolution

## Abstract

Gene transfer between distinct evolutionary lineages has been recognized as a frequent event occurring between viruses and their hosts. This phenomenon has been studied to some extent in animal and plant viruses, not so much in the case of mycoviruses, for which the evolutionary origins of their proteins remain poorly understood. In this study, we have tested the hypothesis of a mosaic origin for mycoviruses’ genomes, with the RNA-dependent RNA-polymerase (RdRp) being of viral origin and the coat protein (CP) resulting from one or more transfer events from the host genome. Firstly, phylogenetic trees were constructed for the RdRps and the CPs from a selection of viruses to address for possible incongruent evolutionary histories. Moreover, a PSI-BLAST search using the CP sequences from the different mycovirus groups retrieved hypothetical proteins (HP) with many orthologues in fungal genomes showing significant sequence homology with the CP from the members within the *Curvulaviridae* family. The structures of these HPs, predicted *in silico* using AlphaFold, tend to show high similarity with viral CPs suggesting the occurrence of gene transfer between viruses and fungi, although no clear function has been yet attributed to these genes in the host. Phylogenetic analyses suggest that this gene transfer could have occurred in multiple independent events. Additional selection analysis supports the notion that the most parsimonious explanation is the transfer of the HP from the host to an ancestral viral genome followed by fast evolution to accommodate the newly acquired protein to function as a CP.

## Introduction

Fungal viruses, or mycoviruses, are widespread in fungi and believed to have evolved at an early stage in the phylogeny of their hosts (Hough et al. 2023). They usually are associated with symptomless infections and are transmitted intra-cellularly during sporogenesis, cell fusion, and cell division (Ghabrial and Suzuki 2008). Their natural host ranges are limited to individuals within the same or closely related vegetative compatibility groups (Ghabrial 1998). Mycoviruses attract considerable attention due to their great diversity in terms of genome organization, replication cycles as well as, their potential to be used as biological control (García-Pedrajas et al. 2019; Sutela et al. 2020; Villan Larios et al. 2023). Until recently, fungal virosphere was believed to be heavily dominated by double-stranded (ds) RNA viruses (Kondo, Botella, and Suzuki 2022). This turned out to be inaccurate as a result of employing biased detection methodologies in the past, since dsRNA was used as a marker for fungal virus infection (Morris and Dodds 1979). Indeed, deep RNA sequencing approaches using small RNA (Donaire and Ayllón 2017; Muñoz-Adalia et al. 2018), dsRNA, or ssRNA inputs (Marzano and Domier 2016; Marzano et al. 2016) for cDNA library construction revealed a large number of (+)ssRNA viruses present in the fungal world. In addition, many (-)ssRNA viruses with non-segmented and segmented genomes have been discovered (Kondo et al. 2013; Liu et al. 2014; Donaire, Pagán and Ayllón 2016; Lin et al. 2019). Moreover, a vast number of unusual viruses with peculiar genome architectures that were previously unknown, have been discovered in fungi (Chiapello et al. 2020; Sutela et al. 2020; Chiba et al. 2021; Forgia et al. 2021; Jia et al. 2021; Linnakoski et al. 2021). These recent studies suggest that the mycovirosphere is dominated by dsRNA and (+)ssRNA viruses, with a more restricted representation of (-)ssRNA and ssDNA viruses. Indeed, neither pararetroviruses nor true dsDNA viruses have yet been reported in fungi.

The genome of encapsidated dsRNA mycoviruses is either segmented or non-segmented and encodes at least an RNA-dependent RNA-polymerase (RdRp) and a coat protein (CP). The RdRp is needed to replicate the dsRNA genome and contains highly conserved motifs across the RNA virosphere. The CPs remain structurally undisturbed throughout the viral cycle and are involved in the organization of the viral genome and the viral polymerase (Luque et al. 2018). Comparative analysis of the amino acid sequences of proteins encoded by dsRNA viruses revealed little similarity between viruses of different genera (Ghabrial 1998). Despite being a highly conserved gene among RNA viruses, phylogenetic analysis of the RdRp did not suggest a monophyletic origin for dsRNA viruses (Koonin 1992). The dsRNA viral RdRps tend to group with different subdivisions of the (+)RNA viruses, suggesting that dsRNA viruses could have originated at multiple events in evolution from different supergroups of (+)RNA viruses (Koonin 1992). Different scenarios were proposed for the emergence of dsRNA mycoviruses members, some involved the acquisition of the CP by the replicative forms (*e.g*., *Totiviridae*) or its loss (*e.g*., *Hypoviridae*) (Ghabrial 1998). Other scenarios involved genome rearrangement and segmentation (*e.g*., *Partitiviridae*) (Ghabrial 1998). Krupovic and Koonin (2017) conducted a systematic comparison of viral proteins with the global database of protein sequences and structures and suggested that viral CPs could have evolved from *bona fide* cellular proteins, in several independent occasions through horizontal gene transfer (HGT).

HGT between viruses and host or virus to virus has been recognized as an important driving force in viral evolution. It was illustrated to occur from cells to viruses (Moreira 2000; Bratke and McLysaght 2008; Moreira and Brochier-Armanet 2008; Moreira and López-García 2009), as well as from viruses to viruses (Koonin and Dolja 2006). Transfer from retroviruses to eukaryotic cells or DNA viruses has also been reported. Transfer from non-retroviral RNA viruses to cells is thought to be extremely rare (Bertsch et al. 2009; Geuking et al. 2009; Mushegian and Elena 2015). Liu et al. (2010) analyzed eukaryotic genome databases for the presence of dsRNA virus-related sequences, and suggested that genes from partitiviruses and totiviruses were likely transferred from viruses to eukaryotic genomes, and some of these genes are functional. However, no HGT event from the host to virus has yet been reported in dsRNA mycoviruses. In this study, we investigated this hypothesis through sequence and structure similarity across RdRps and CPs of the families *Chrysoviridae*, *Curvulaviridae*, *Megabirnaviridae*, *Partitiviridae*, *Totiviridae*, and few extra unclassified dsRNA mycoviruses, looking for eukaryotic protein candidates which could have been hijacked by dsRNA mycoviruses, and we provide evidence that members of the family *Curvulaviridae* had probably acquired their CPs from the fungi.

## Results

### Expanding the mycoviruses sequences used for this study

We retrieved the RdRp and the CP sequences of different mycoviruses using a set of nine species belonging to different genera and families (Table S1). The number of hits generated by PSI-BLAST for the CP ranged from 16 hits for Beauveria bassiana non-segmented RNA virus 1 (BBRV1) to 229 hits for Heterobasidion partitivirus 1 (HPV1). In the case of the RdRp, the number of hits obtained was 476 for Trichoderma harzianum bipartite mycovirus 1 (ThBMV1) (Liu et al. 2019), 492 for BBRV1 and 500 for the other viruses exceeding in all the cases the hits obtained by the CP. After filtering the PSI-BLAST hits, we ended up with 251 virus sequences for the analysis for each of the proteins (Table S2). Since most viruses were only classified at family level (Table S2), classified species were used to assign unclassified ones when clustering together in phylogenies for the rest of the study.

### Phylogenetic congruence between the RdRp and the CP proteins of mycoviruses

To examine whether the CP and the RdRp of dsRNA mycoviruses shared the same evolutionary history, we produced a phylogenetic tree for each of the two proteins and compared them through co-phylogenetic analysis (Fig. 1). Both trees showed five clades, a major clade comprised species from the *Partitiviridae* family, which constituted the majority of the sequences from our study, as well as species assigned to the *Curvulaviridae* and *Totiviridae* families (Fig. 1). The partiti-like clades comprised three sub-clades grouping members belonging to alpha-, beta- and gammapartitiviruses (Fig. 1). Two other separated clades comprised alpha- and betachrysoviruses belonging to the *Chrysoviridae* family. An additional clade comprised members of the *Megabirnaviridae* family and a last one comprised unclassified species. The co-phylogenetic tree of the RdRp *vs* the CP showed incongruence within groups of each clade (at the genus level), but also between the different clades (family level) (*e.g*., *Betachrysovirus* and *Megabirnavirus*) (Fig. 1). This incongruence may either suggest that the RdRp and the CP might have a different evolutionary history, or recombination may have contributed to the discrepancy observed between these two proteins. To verify the latter hypothesis, recombination analysis was carried out by concatenating the RdRp and the CP genes; 17 breakpoints were identified (Table S3, Fig. S1), in most of the cases the breakpoint seemed to occur either in the RdRp gene or in the CP gene (Fig. S1), except in the case of Plasmopara viticola lesion associated partitiviruses 3 and 10, where the breakpoint lied between these two genes (Fig. S1). Moreover, no recombination was often detected in viruses for which the RdRp and the CP positioned differently within the same clade (*e.g*., Rhizoctonia solani partitivirus 8, and Plasmopara viticola lesion associated partitivirus 4) or between different clades (*e.g*., *Betachrysovirus* and *Megabirnavirus*). Overall, co-phylogenetic analysis shows that the discrepancies between the RdRp and the CP of the mycoviruses were in few cases explained by recombination.

**Fig. 1.**
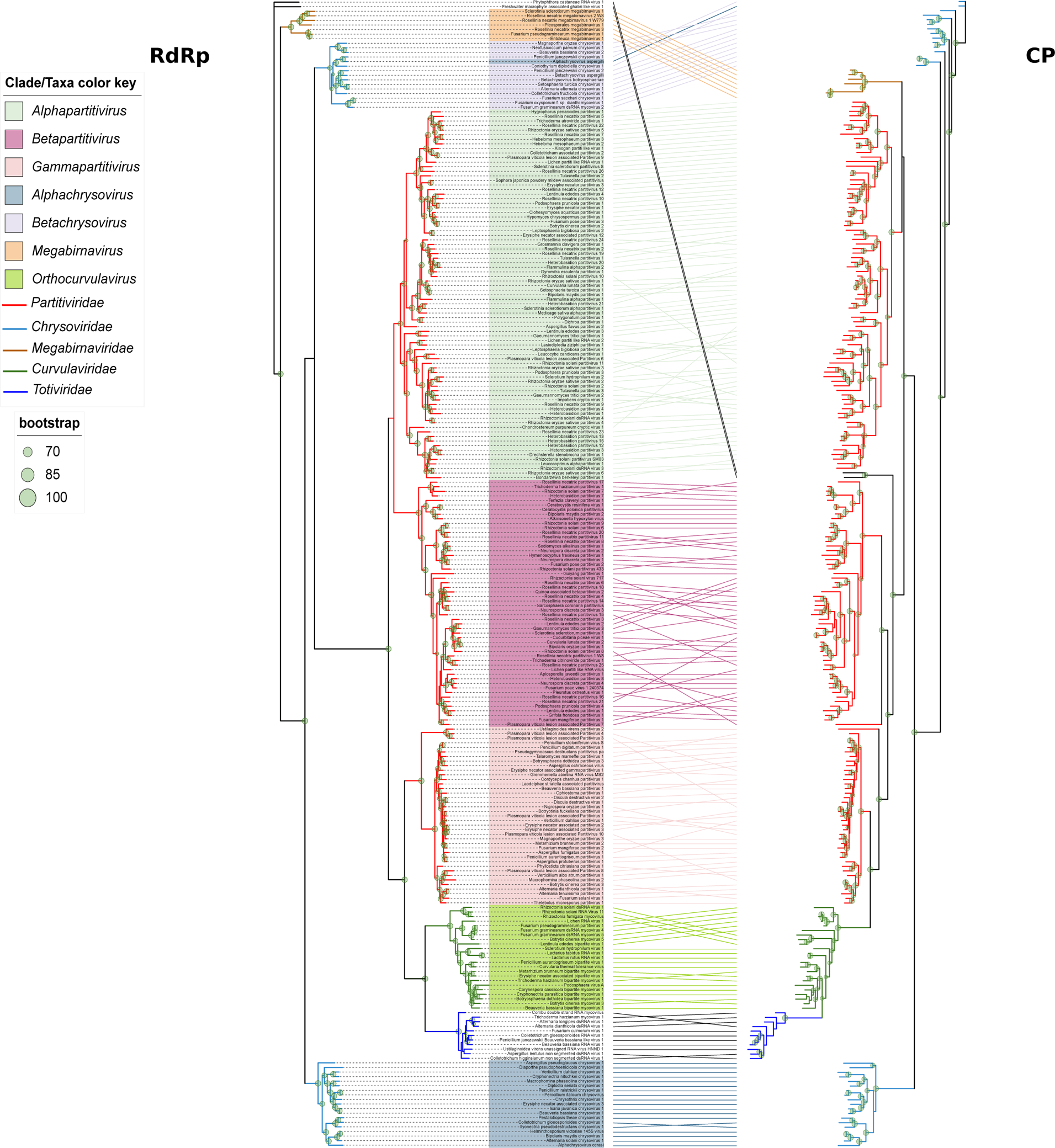
Co-phylogenetic tree showing the phylogenetic relationships between the RNA-dependent RNA-polymerase (RdRp, right), and the coat protein (CP, left) of mycoviruses.

### Relationships between the CPs of mycoviruses

We then investigated whether the CPs might share a common ancestor. We generated and compared the tertiary structure of 64 random CPs from the different families of dsRNA mycoviruses. We predicted the presence of at least three different CP structures (*z* < 2 in DALI) (Fig. 2A, Fig. 2C). The partiti-like CP group (Fig. 2A), which comprised the members from the family *Partitiviridae* (Fig. 2A, Fig. 2B), is divided into three subgroups representing each of the genera alpha-, beta- and gammapartitiviruses, but still, they are significantly similar in comparison to the other CPs. The curvula-like CP (Fig. 2A), which comprised members from the *Curvulaviridae* and the *Totiviridae* families sharing similar motifs, and a third group comprised members from the *Chrysoviridae* family which was also divided in two subgroups (alpha- and betachrysoviruses). The dendrograms based on the CP structures showed different CP-like groups that are clearly different, suggesting that the CPs of mycoviruses are polyphyletic.

**Fig. 2.**
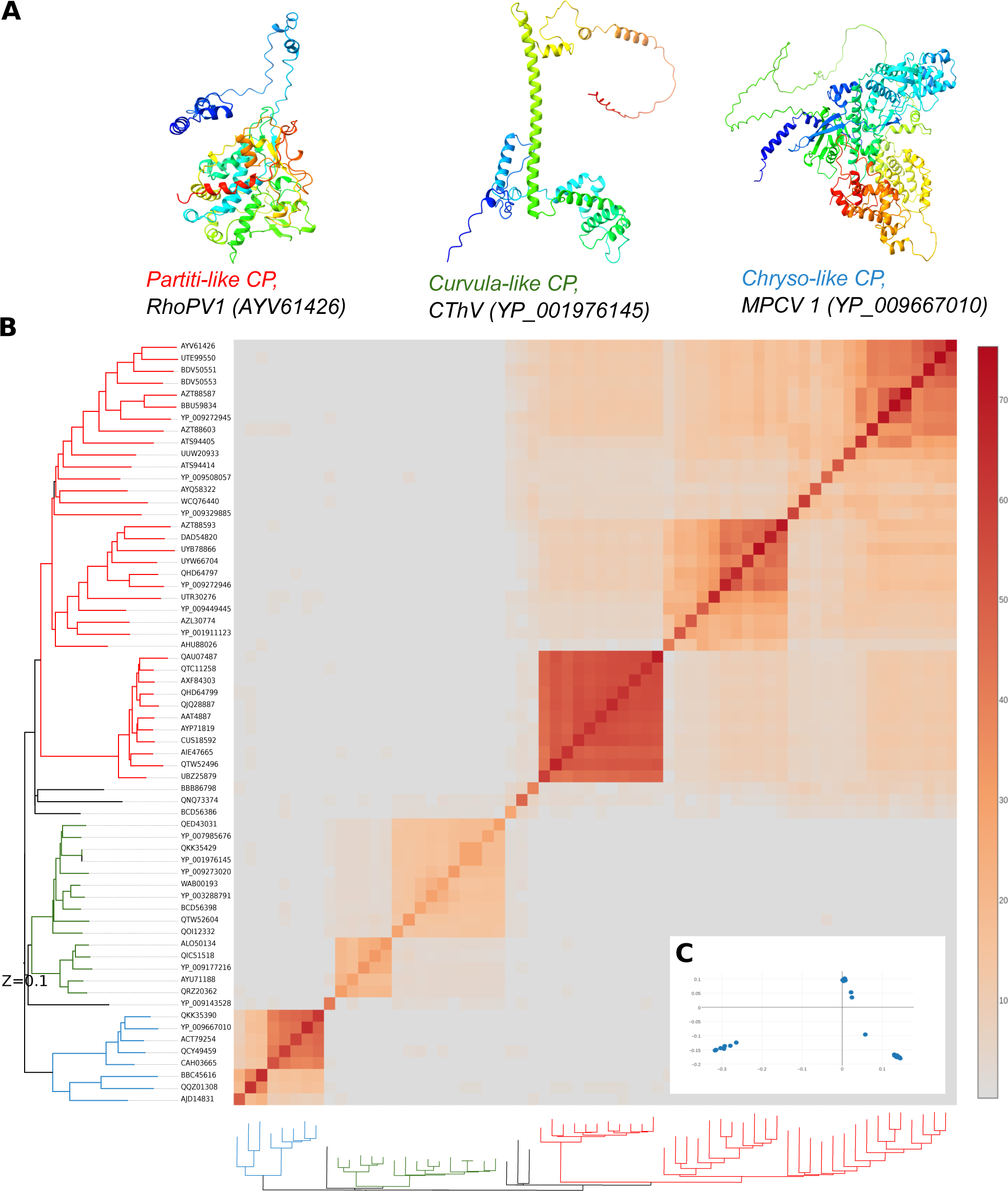
Viral proteins. (A) A selection of representative coat protein structures of mycoviruses. All structures are colored using the rainbow scheme from blue (N terminus) to red (C terminus). (B) Relationships between the different viral proteins. The matrix and cluster dendrograms are based on the pairwise X score comparisons using DALI. Virus proteins are colored red for partiti-like CPs, green for curvula-like CPs and blue for chryso-like CPs in the dendrogram. (C) Principal component analysis of the different viral proteins.

### Cellular orthologs for the CP genes in the host

To further test the hypothesis of HGT from the host as the potential origin of the CPs, we sought orthologous proteins from the host by conducting PSI-BLAST searches using the same viral species above as a query. The number of fungal hits obtained ranged from 0 for Fusarium solani virus 1 and Fusarium pseudograminearum megabirnavirus 1 to 335 to Coniothyrium diplodiella chrysovirus 1 (Table S1). After filtering, we ended up with 279 fungal proteins used for the analysis (Table S4). Next, we examined the phylogenetic relationships between the CPs and the fungal proteins (Fig. 3). Two clades were formed. The first clade comprised only CPs of viruses from the *Partitiviridae*, *Chrysoviridae* and *Megabirnaviridae* families (Fig. 3). The second clade comprised the host proteins HP as well as curvula-like and toti-like CPs, as well as some divergent viral lineages from partitiviruses (Ustilaginoidae virens partitivirus 2 and Rhizoctonia solani partitivirus 8) and from an unclassified virus (Phytophtora castaneae RNA virus 1) (Fig. 3). In the case of the curvula-like CPs, the virus proteins clustered within the fungal HPs in seven different clusters (Fig. 3). In contrast, the toti-like CPs formed a unique, divergent cluster within the host proteins.

**Fig. 3.**
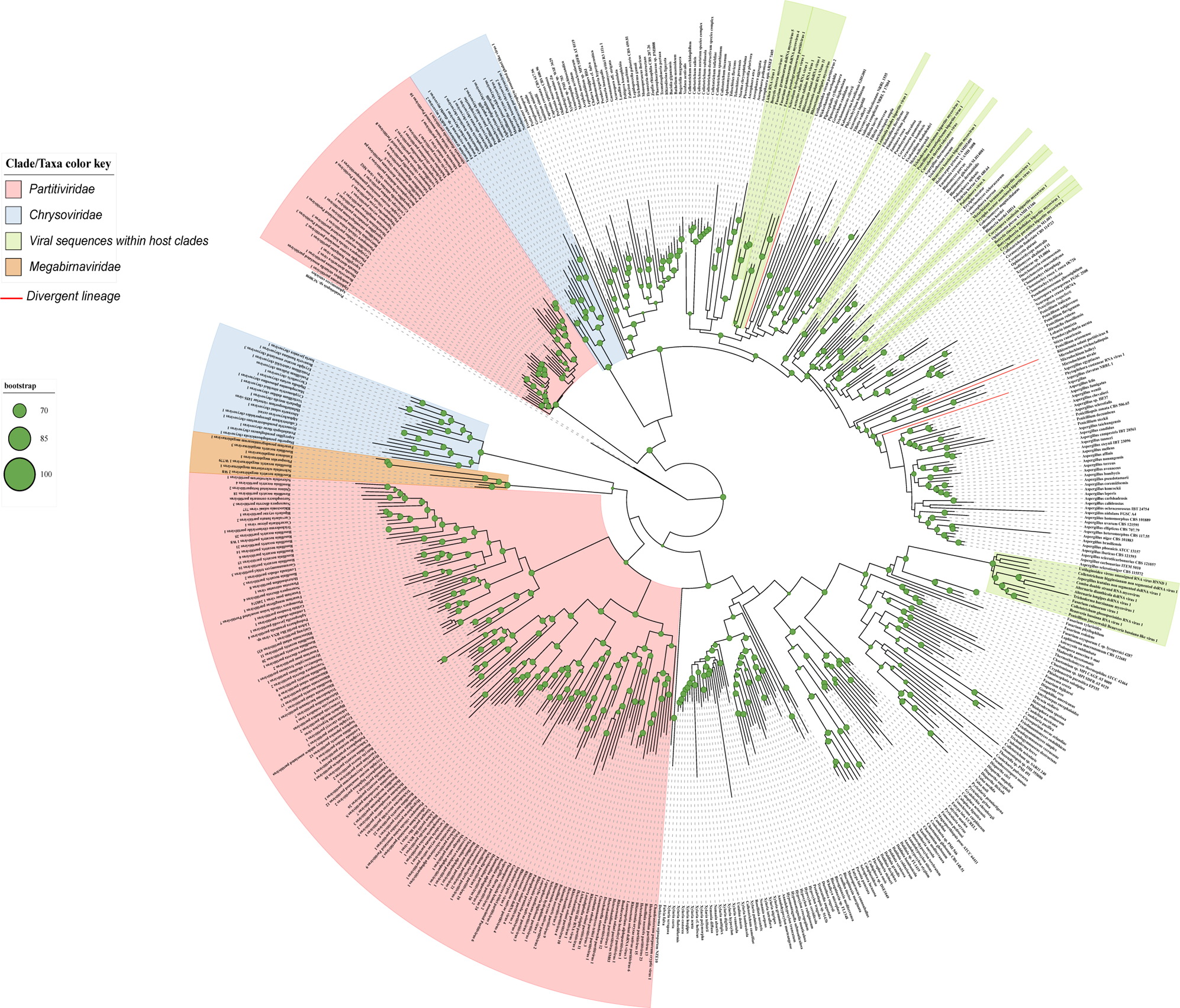
Phylogenetic tree of viral coat proteins and their related sequences from the host.

### Relationships between the virus CPs and its orthologs from the host

The relationships among these proteins were further explored by comparing the tertiary structure of the viruses and the fungal proteins that clustered closer in the phylogenetic tree. A similar folding was observed in the case of *Aspergillus coremiformis*, and *Ascosphaera pollenicola* in comparison to curvula-like CPs (Fig. 4A and 4B). Other HPs like *Colletotrichum sublineola* and *Emmonsia crescens* showed similar structure but with additional folding in the long alpha helices (Fig. 4A and 4B). Structure prediction of the sequences that cluster closer to toti-like CPs, was either poor (Fig. S2A) or was dissimilar to toti-like sequences proteins even when trying different structure alignment settings (Fig. S2B). The distance matrix and its corresponding dendrograms (Fig. 4C) further confirmed the presence of a significant similarity between the curvula-like virus protein and host proteins. Despite the significant similarity found using DALI (*z* > 2), the 3D structure of the toti-like and curvula-like virus seems to be different, suggesting that the DALI score values observed may be due to some shared motifs (alpha helices). Most of the virus CP orthologs from the host were assigned either as hypothetical proteins or proteins of unknown functions (Fig. 4B) except in the case of *Ascosphaera pollenicola*, from which the curvulavirus CP homolog is annotated as peroxisomal membrane receptor (PTS1) (Fig. 4B). Overall, our analysis suggests that the curvula-like CP shares a common evolutionary history with fungal HPs, whose comparison at structural level reflects the occurrence of HGT between the host and the viruses. Structural similarity between the fungal proteins and curvula-like proteins, suggests that fungal proteins are capable of forming assemblies resembling virus-like particles (Fig. 4A and 4B).

**Fig. 4.**
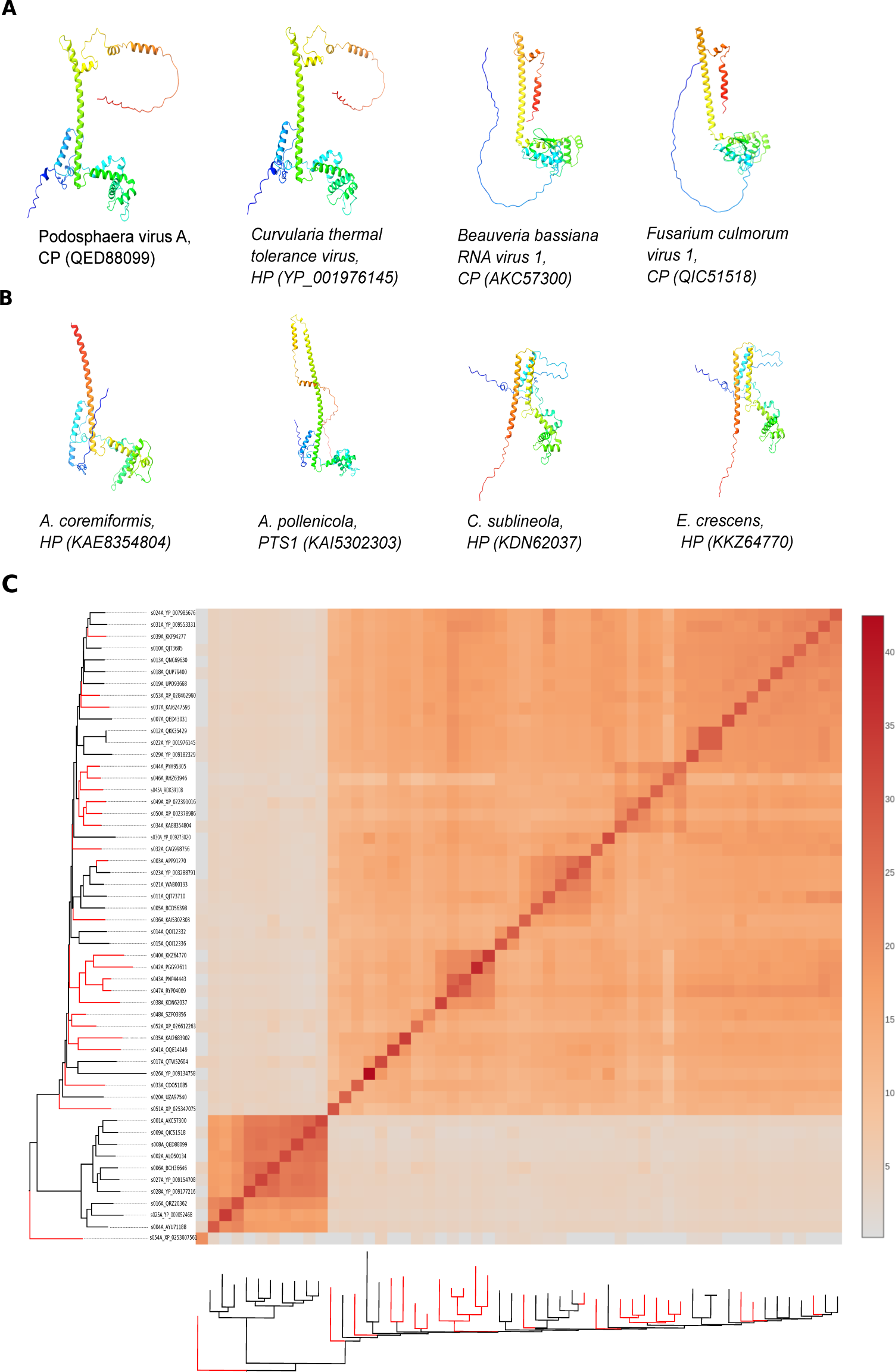
Viral CPs and their orthologous cellular proteins. (A) A selection of viral curvula-like CP structures. (B) A selection of host proteins similar to curvula-like CPs. All structures are colored using the rainbow scheme from blue (N terminus) to red (C terminus). (C) Relationships between cellular and viral proteins. The matrix and cluster dendrograms are based on the pairwise *Z* score comparisons calculated using DALI. Host proteins are colored red in the dendrogram. HP: Hypothetical protein, PTS1: peroxisomal membrane signal receptor.

A possible concern of these analyses is the presence of one or more viruses infecting the fungi that were sequenced and thus the HP being actually CP proteins from unknown viral species. To rule out this possibility, we ran PSI-BLAST using the CPs and the RdRps from curvula-like viruses against those fungi genomes which showed similarity with viral proteins. In all cases, we got hits for the CPs, but none for the RdRps, pointing that HPs are likely host proteins and not from other viruses present in the cells (data not shown).

### Directionality of the gene transfer

Next, we sought to explore the directionality of the HGT. To do so, we focused on the curvula-like CPs since it showed significant structure similarity with the host HPs. We compared the relative evolutionary rate of the 23 curvula-like CPs to 34 partiti- and chryso-like CPs (including both alpha- and betachrysoviruses) to their RdRps by calculating nonsynonymous (*d_N_*) to synonymous (*d_S_*) substitutions ratios (*ω*) at the tree branches, and tips levels (Table S5 and Table S6). Since the RdRp shows differences in topology compared to the CP and to make the comparisons possible at the branch levels, the evolutionary rate was calculated on maximum-likelihood “species trees” generated by concatenating the RdRp and the CP sequences (Fig. S3 and S4). The evolutionary rate of the CP from curvula-like viruses (HP) was significantly higher when compared to the partiti-like CPs (CP1) at branches level (8.19·10^13^ *vs* 2.86·10^7^, *P* < 0.01), at tips level (21.32 *vs* 14.05, *P* < 0.01), as well as after combining tips and branches (3.59·10^13^ *vs* 1.29·10^7^, *P* < 0.01) (Fig. 5A). When compared to the chrysolike-CPs (CP2), the HP seems to evolve faster than the CP2 at branches (1.10·10 ^11^ *vs* 2.17·10^10^) and tips (29.44 *vs* 3.97), but at a similar rate when combining both tips and branches (1.27·10^10^ *vs* 1.10·10^10^). However, the differences noticed at branch and tip levels were not significant (*P* > 0.05). Our analysis showed that the CP of curvulaviruses and chrysoviruses are evolving at a similar rate, but still faster than the partiti-like CPs.

**Fig. 5.**
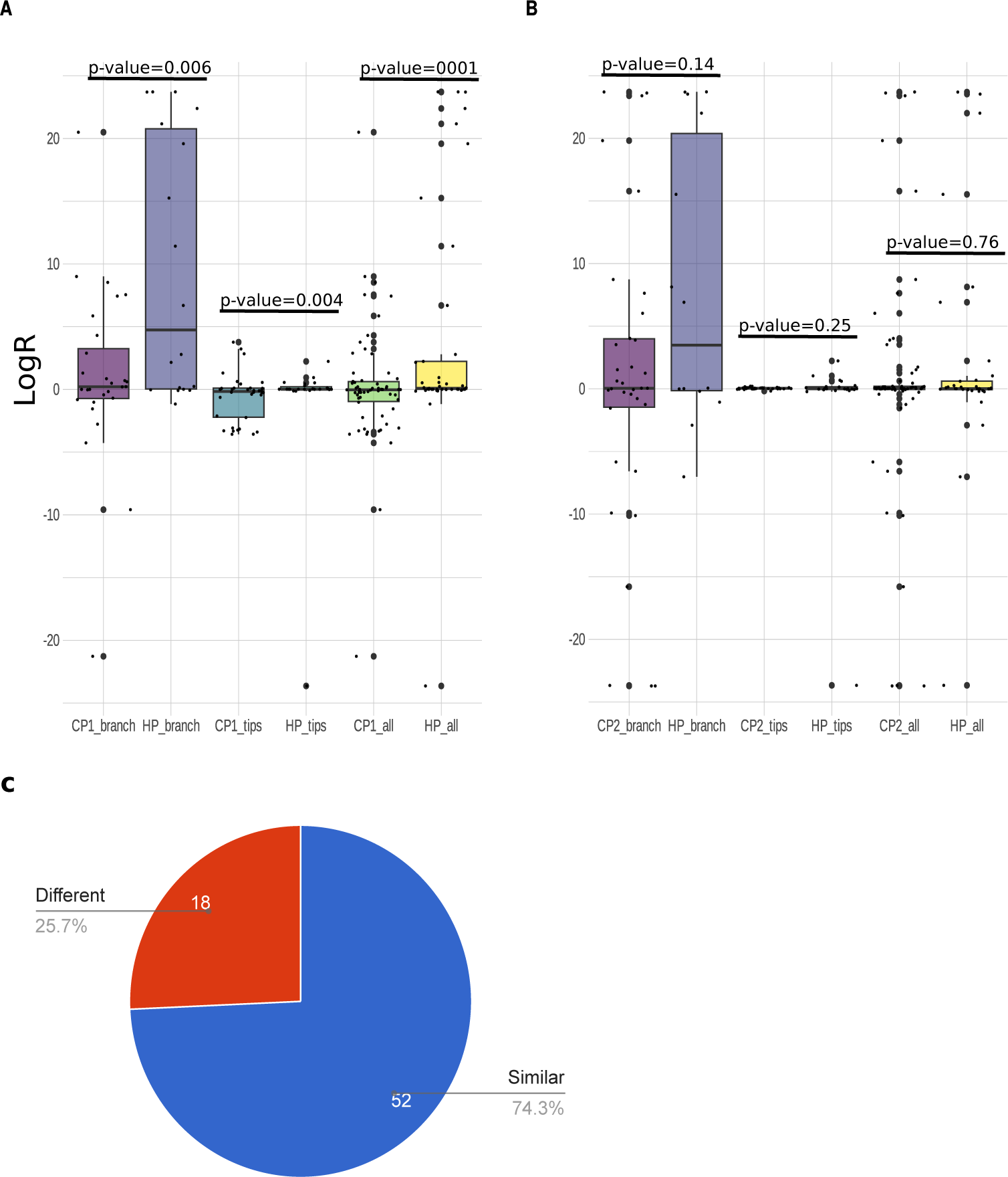
Evolutionary rates comparisons. (A) Comparison of evolutionary rate of the curvula-like CP (HP) with partiti-like CP (CP1). (B) Comparison of evolutionary rate of curvula-like CP (HP) with chryso-like CP (CP2). The comparison was made at tip, branch and combining both tips and branches. (C) Comparison of the evolutionary rate of curvula-like CP orthologous from the host with other 70 random proteins.

We then investigated the behavior of the curvula-like CP homologs in the host genomes. We first selected 70 random proteins from the fungus *Penicillium roqueforti* (Table S7), then we retrieved the homologs of these proteins from 24 fungi (Table S8, Table S9). We calculated the *ω* ratios at the branch and tips levels (Table S10). Because of the differences between the phylogenetic tree of each of the proteins, and to be able to compare the different branches, the *ω* was calculated on a maximum-likelihood species tree, generated by concatenating all 71 proteins together (Fig. S5). The evolutionary rate of the curvula-like HP homologs from the host was compared to the 70 randomly selected proteins. In 74.3% of the cases (52/74) (Fig. 5C), the evolutionary rate of the curvula-like CPs was like the host proteins rate, and it evolved differently when compared to the other 18 proteins (25.7%, 18/74). The curvula-like CPs tend to evolve much faster than these 18 host proteins (Table 1).

**Table 1.** List of proteins evolving slower than the curvulavirus CP orthologs in the fungi genome and their functions.

| Protein ID | Description | $\omega_{HP}/\omega_{FP}^1$ | Role |
| --- | --- | --- | --- |
| CDM27110 | Calpain-like protease palB/RIM13 | $1.03 \cdot 10^{11}$ | Required for the proteolytic cleavage of the transcription factor RIM101 in response to alkaline ambient pH. |
| CDM32354 | Eukaryotic translation initiation factor 2A | $1.03 \cdot 10^{11}$ | Functions in the early steps of protein synthesis of a small number of specific mRNAs. Acts by directing the binding of methionyl-tRNA<sub>i</sub> to 40S ribosomal subunits |
| CDM34807 | Diphthine synthase | $4.37 \cdot 10^{10}$ | Amidase that catalyzes the last step of diphthamide biosynthesis using ammonium and ATP. Diphthamide biosynthesis consists in the conversion of an L-histidine residue in the translation elongation factor eEF-2 (EFT1 or EFT2) to diphthamide |
| CDM33363 | Probable T-complex protein 1 subunit eta | $5.39 \cdot 10^{10}$ | Molecular chaperone; assists the folding of proteins upon ATP hydrolysis |
| CDM33016 | Peroxisomal targeting signal receptor | $9.08 \cdot 10^{10}$ | Receptor required for the peroxisomal import of proteins containing a C-terminal PTS2-type peroxisomal targeting signal, such as 3-oxoacyl-CoA thiolase |
| CDM33059 | COP9 signalosome complex subunit 4 | $2.75 \cdot 10^{10}$ | Complex involved in various cellular and developmental processes. The CSN complex is an essential regulator of the ubiquitin (Ubl) conjugation pathway by mediating the deneddylation of the cullin subunits of the SCF-type E3 ligase complexes |
| CDM31535 | Alcohol dehydrogenase 1 | $5.95 \cdot 10^{10}$ | Oxidizing mainly primary and secondary aliphatic alcohols, utilizing NAD<sup>+</sup> as a cosubstrate |
| CDM32496 | Pre-mRNA-splicing factor brr2 | $2.38 \cdot 10^{10}$ | Required for pre-spliceosome formation, which is the first step of pre-mRNA splicing |
| CDM37865 | Sec63 domain | $4.29 \cdot 10^{10}$ | Acts as component of the Sec62/63 complex which is involved in SRP-independent post-translational translocation across the endoplasmic reticulum (ER) and functions together with the Sec61 complex and KAR2 in a channel-forming translocon complex |
| CDM35843 | Signal recognition particle 54 kDa protein homolog | $5.69 \cdot 10^{10}$ | SRP is required for the cotranslational protein translocation for ER import and preferentially recognizes strongly hydrophobic signal sequences. It is involved in targeting the nascent chain-ribosome (RNC) complex to the ER |
| CDM26399 | Protein vts1 | $8.29 \cdot 10^9$ | RNA-binding protein involved in post-transcriptional regulation through transcript degradation of SRE (SMG-recognition elements) bearing mRNAs |
| CDM28512 | TRAPP II complex, TRAPPC10 | $1.81 \cdot 10^{10}$ | Required for both the cytoplasm-to-vacuole targeting (Cvt) pathway and starvation-induced autophagy through its role in ATG8 and ATG9 trafficking |
| CDM32296 | Mediator of RNA polymerase II transcription subunit 12 | $5.03 \cdot 10^{10}$ | Coactivator involved in the regulated transcription of nearly all RNA polymerase II-dependent genes. |
| CDM32001 | Threonine synthase | $5.85 \cdot 10^9$ | Catalyzes the gamma-elimination of phosphate from L-phosphohomoserine and the beta-addition of water to produce L-threonine. |
| CDM33552 | Serine/threonine-protein kinase srk1 | $2.52 \cdot 10^{10}$ | During osmotic stress, inhibits the G2/M transition in a sty1 stress-activated MAPK pathway-dependent manner |
| CDM33998 | Aspartyl-tRNA synthetase, cytoplasmic | $8.28 \cdot 10^9$ | Catalyzes the attachment of L-aspartate to tRNA(Asp) |
| CDM36914 | Anthranilate synthase component 1 | $9.07 \cdot 10^9$ | Component I catalyzes the formation of anthranilate using ammonia rather than glutamine, whereas component II provides glutamine amidotransferase activity. |
| CDM31360 | Peptidyl-prolyl cis-trans isomerase-like 3 | $4.91 \cdot 10^9$ | PPIases accelerate the folding of proteins. It catalyzes the cis-trans isomerization of proline imidic peptide bonds in oligopeptides. |

<sup>1</sup>HP, viral ortholog proteins from the host; FP, fungal proteins.

## Discussion

Under the assumption that the RdRps and the CPs of dsRNA mycoviruses shared the same evolutionary history since virus emergence, their phylogenies should be topologically congruent (Göker et al. 2011). However, a combination of events such as recombination, host-virus gene interactions or gene transfer could lead to the topological incongruence observed in our analysis. In fact, these types of discrepancies were also reported from different studies (Roossinck 2019). In partitiviruses, for instance, phylogenetic analysis of the RdRp showed separate groups; some are specific to plants or to fungi (Li et al. 2009; Roossinck 2010; Nibert et al. 2014). However, the CP genes group closely to their host than the RdRp genes (Roossinck 2019). Investigating the relationships among the CPs of different dsRNA mycovirus groups revealed clear differences in the folds for CPs belonging to different families suggesting that the CPs of dsRNA mycoviruses are not monophyletic. The observations from our study, and from previous ones, provide compelling evidences that the RdRp core gene acquired the CP genes at various times and probably from different sources.

This stands in contrast to the case of viral RdRps, which have no closely related homologs in known cellular organisms (Koonin and Dolja 2006; Kazlauskas et al. 2016). Recent studies on the origin of viral capsid proteins revealed unexpected similarities in the folds of CPs from viruses infecting hosts from different cellular domains, testifying to the antiquity of the CPs and the evolutionary connections between the viruses that encode them (Krupovic and Bamford 2008; Kormelink et al. 2011; Abrescia et al. 2012). Searching for CPs of dsRNA mycovirus orthologs in the host genomes using PSI-BLAST, coupled with phylogenetic analysis, yielded CPs from totiviruses and curvulaviruses which tightly clustered within host proteins. On the one hand, the CPs of totiviruses constituted a divergent cluster within the host proteins suggesting that these proteins have been originated from a single event, although, no clear similarity was observed when comparing fungi and totivirus proteins at the structural level. On the other hand, the CPs from curvulaviruses grouped with fungal proteins in different clades. This grouping indicates that the curvulavirus CPs share a similar evolutionary history with fungal proteins and one of the proteins has originated at multiple events in evolution from the other protein groups. In contrast to totivirus CPs, the CPs from curvulaviruses showed a quite similar folding structure with the host proteins.

The *Curvulaviridae* family was created recently after the discovery of Curvularia thermal tolerance virus (CThTV) (Márquez et al. 2007). Their bisegmented genome encodes an RdRp, and a protein annotated in most of the species belonging to this family as hypothetical protein (Walker et al. 2021). Nevertheless, the identification of spherical virions of 26-29 nm with CThTV infection suggests that these hypothetical proteins are coat proteins (Márquez et al. 2007).

The majority of the fungal species hosting the curvula-like orthologs belonged to the *Ascomycota* division. Exceptions are *Acaromyces ingoldii* and *Pseudomicrostroma glucosophilum*, which belonged to the *Basidiomycota* division probably because the former division is considerably richer in number of species than the latter (Wang et al. 2010), but this would also suggest that curvulaviruses have evolved at a very early stage in the phylogeny of the ascomycetes, and there have been few events where dsRNA mycoviruses may have moved across fungal lineages. Most fungal genomes with these proteins are either poorly annotated or in a draft stage. Due to these pitfalls in genome annotation, most proteins are assigned either as hypothetical proteins or proteins of unknown function, making it unfeasible to assign a function to these proteins in host. The only exception was the case of *A. pollenicola*, from which the curvulavirus CP ortholog is annotated as peroxisomal membrane receptor (PTS1). This protein seems to bind to cargo proteins containing a PTS1 peroxisomal targeting signal in the cytosol and translocate them into the peroxisome matrix by passing through the PEX13-PEX14 docking complex along with cargo proteins (Skowyra and Rapoport 2022). Still, it is possible that this fungal protein was incorrectly annotated.

Phylogenetic relationships and structural analyses between the curvulaviruses CPs and their homologs from the fungi illustrate a clear case of gene transfer between fungi and viruses. One of the challenges was to determine the directionality of that transfer. One hypothesis we sought was if the virus acquired its CP from the host. This being the case, the relative evolutionary rate of the curvulavirus CPs when compared to their RdRps would be higher than the one of the CPs from other viruses, reflecting a recent acquisition followed by a (still undergoing) process of adaptation. The curvulavirus CPs tend to evolve faster than the partitivirus CPs but at a similar rate than chrysovirus CPs. This may suggest that these viruses acquired their CPs in a recent time in comparison to partitiviruses, although, no homologs of chrysoviruses were recovered from the fungi, which suggest that their CPs were probably acquired from other organisms and went through gene duplication (Luque et al. 2010). Rappoport and Linial (2012) suggested that along viral evolution in metazoan, the host-originated sequences accommodate simplified domain compositions, and are significantly shorter than their metazoan counterparts. However, the lengths of the curvulavirus CPs and their homologs tend to be of a comparable size.

According to our hypothesis, we expect host HPs to evolve at the same rate than other host genes, while if the direction of HGT was from viruses to hosts, then HPs should evolve faster than the average host gene. The comparison of the evolutionary rate between HPs and a set of 70 fungal proteins showed that in ∼75% of the cases the CPs homologs evolved similar than host proteins, while in ∼25% evolved faster. After investigating the role of the set of slow-evolving proteins, we found that they were enriched in housekeeping and essential genes, which are expected to evolve particularly slow.

The occurrence of gene transfer from dsRNA mycoviruses and their host was already reported. Liu et al. (2010) concluded that viral homologs integrated eukaryotic genomes and some of these transferred genes are conserved and expressed in eukaryotic organisms, what suggests that these viral genes are also functional in the recipient genomes. In the case of curvulaviruses, our analysis-based on the behavior of the CPs and their homologs in the host-provides evidence for a gene transfer from the host to the virus raising the possibility of two different scenarios for its occurrence (Fig. 6). In the first one, curvulaviruses in their early stage of emergence were naked dsRNA replicative forms and through their evolution with the host they hijacked their CPs from the host (Fig. 6A). The second scenario involves that curvulaviruses had an ancestral CP which was lost and/or substituted by a host protein through their co-evolution with the host (Fig. 6B).

**Fig. 6.**
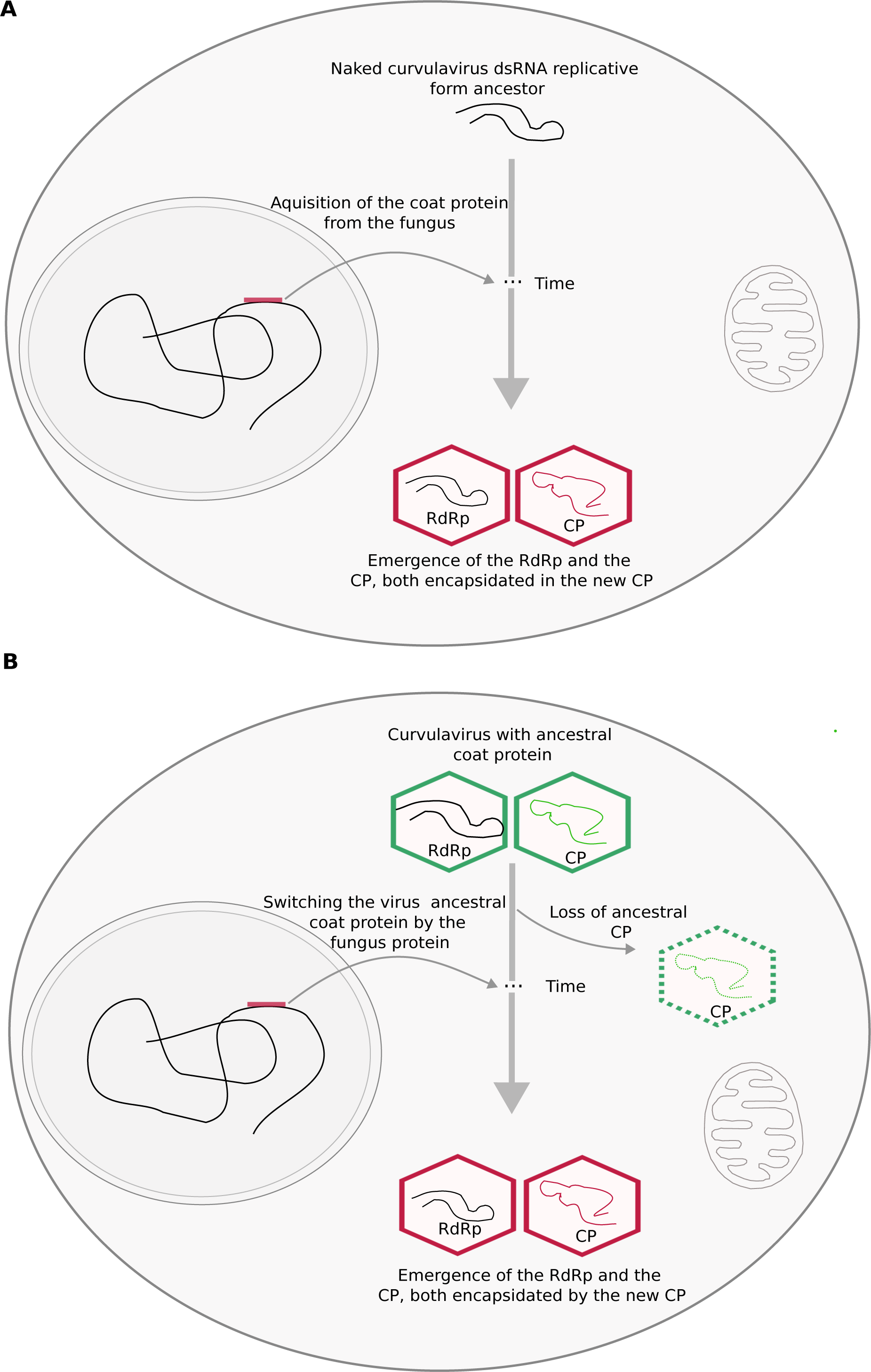
Scenarios for the emergence of members from the *Curvulaviridae* family. (A) Curvulaviruses were naked replicative forms and through their evolution they required the coat protein from the fungi. (B) Curvulaviruses had an ancestral coat protein which was switched by the fungi proteins. RdRp, RNA dependent RNA polymerase; CP, coat protein.

## Material and methods

### PSI-BLAST searches for retrieving virus and host protein sequences

Amino acid sequences for the RdRp, CP and putative CP (pCP) from six species, representing mycoviruses’ genera and families (Table S1), were used to extend the number of viral sequences from each genus or family for the analysis. For each virus, sequence-similarity searches were performed using PSI-BLAST algorithm (Altschul et al. 1990) against the non-redundant protein sequence database at the National Center for Biotechnology Information (NCBI) (https://blast.ncbi.nlm.nih.gov/Blast.cgi) using three iterations with default algorithm parameters to retrieve viral sequences. For distant sequence similarity detection (sequences from the fungi), PSI-BLASTP was run using experimental database option with five iterations, selecting only fungi through the iterations. Amino acid sequences and their corresponding nucleotide coding sequences were downloaded using BATCH ENTREZ (https://www.ncbi.nlm.nih.gov/sites/batchentrez). Viral sequences were analyzed by using the sequence demarcation tool (SDT) version 1.2 program (Muhire, Varsani and Martin 2014) and filtered based on the following criteria (*i*) species with partial sequences for both the CP and the RdRp were removed (*ii*) species that had one partial sequence protein, were substituted by a complete sequence when possible, otherwise removed (*iii*) species showing more than 96% of identity for the RdRp and the CP at amino acid level were considered as the same species, thus only one was selected for the analysis, (*iv*) species reported to infect plants were discarded from the analysis. In the case of host proteins, when hits recovered from multiple strains of the same species, only one hit was selected for the analysis, and sequences annotated as partial were removed. The recovered virus species were further classified based on the Taxonomy browser from NCBI (https://www.ncbi.nlm.nih.gov/taxonomy).

### Alignments, recombination analysis, and phylogenetic trees

Sequences were aligned using MUSCLE version 5 by choosing the-super5 option (Edgar 2022) and were verified manually. Phylogenetic trees were constructed by using the maximum-likelihood (ML) method using IQ-TREE version 2.2.0 (Minh et al. 2020). The best amino acid substitution model for each set of multiple sequence alignment (MSA) was determined by using ModelFinder (Kalyaanamoorthy et al. 2017). VT+F+I+I+R9 model was used for the MSA of the RdRps, VT+F+I+I+R7 model was used for the MSA of the CPs, Q.pfam+F+R7 and Q.pfam+F+R6 models were used for the MSA of the concatenated RdRps and CPs respectively, VT-R9 was used for the MSA of the CPs and fungal proteins and Q.yeast+F+I+I+R5 was used for the MSA of the concatenated sequences from the set of 70 host proteins. Amino acid sequences were concatenated using the seqkit toolkit by choosing the concat option (Shen et al. 2016). PAL2NAL program (Suyama, Torrents and Bork 2006) was used to convert MSA of proteins and the corresponding DNA sequences into a codon alignment. Putative recombination sites were evaluated using the RDP5 program (Martin et al. 2021) by using the following methods: RDP, GENECONV, Maximumx2 (MaxChi), Chimera, BootScan, SisterScan (SiScan), and 3Seq with default settings. Recombination events were considered significant if they were detected by at least five methods. The co-phylogenetic tree was produced using the cophlyo function from the phytools package in R by setting the rotate.multi argument to TRUE. The generated phylogenetic trees were visualized and annotated using R script and ITOL version 6 (Letunic and Bork 2007).

### Structure analysis

Structures of selected proteins from viruses and fungi were predicted using AlphaFold2 (Jumper et al. 2021) run online through “ColabFold version 1.5.2” (Mirdita et al. 2022). Structural similarity matrices from all-against-all structure comparisons as well as corresponding dendrograms were obtained using the latest release of the DALI server (Holm 2022). To investigate structure similarity between virus and fungi proteins, structures were aligned using the MatchMaker algorithm implemented in UCSF Chimera (Pettersen et al. 2004) by choosing the best-aligning pair of chains between reference and matching structures and by using Needleman-Wunsch algorithm. Structures were visualized using the same software. Structural similarities between viral proteins and between fungal and viral proteins were evaluated based on the DALI *z* score, which is a measure of the quality of the structural alignment. Values of *z* > 2 (*i.e.*, ±2 SD above expected by chance) are usually considered significantly similar (Holm et al. 2023).

### Searching for common proteins among different fungi and for viral sequences within fungi

Fungal genomes, from which proteins showed similarity with viral proteins, were downloaded from the EnsemblFungi database (http://fungi.ensembl.org/index.html), except for *Podoshpaera aphanis* and *Trichotecium roseum* which were downloaded from NCBI (Table S2). A protein database for each of the fungi was built using the makeblastdb command. The percentage of annotated proteins from each genome was obtained by dividing the number of proteins annotated as non-hypothetical by the total number of the proteins (Table S8). The well-annotated *Penicillium roqueforti* genome (Table S8) was selected to retrieve 70 random non-redundant proteins (Table S7). A fasta file containing these protein sequences was used to find orthologous proteins from other fungi. Orthologs were obtained by using the BLASTp algorithm against each fungus database allowing only one target sequence for each protein and setting the *E*-value £ 0.001. To search for viral sequences within fungi genomes, fasta files containing the RdRps and the CPs from curvula-like viruses were used for PSI-BLAST search against each fungus database using five iterations.

### Evolutionary rate calculation

Adaptive branch-site random effects model (aBSREL) was used to calculate the nonsynonymous (*d_N_*) to synonymous (*d_S_*) substitutions ratio *ω* = *d_N_*/*d_S_* at the branch level and to determine the ones under positive selection. aBSREL analysis were conducted through HyPhy version 2.5 (Kosakovsky Pond et al., 2005) by using the MSA at codon levels and their corresponding phylogenetic trees as an input. The branch lengths and nucleotide substitution biases were obtained under the nucleotide GTR model. The branches with zero lengths were removed from the analysis. When evaluating evolutionary rates associated to HGT events from the host to the virus, a relative evolutionary rate was calculated as the ratio of the *ω*_CP_ to *ω*_RdRp_, *R_virus_* = *ω*_CP_/*ω*_RdRp_. The distributions of *R_virus_* values, obtained from different CPs (or pCPs), were first tested for normality using Shapiro-Wilk test and further compared using the non-parametric paired samples Wilcoxon test run in R using the shapiro.test() and wilcox.test() functions, respectively. When evaluating the alternative hypothesis of transfer from the virus to the host, the *R_host_*= *ω*_HP_/*ω*_FP_ ratio was obtained by dividing the *ω* of the CP ortholog proteins (HP) from the host by the *ω* of each of the randomly selected fungal proteins (FP) and compared as above. For graphic representation, *R* values were transformed using the asinh transformation (Johnson and Krishnan 2022).

## Supplementary data

**Fig. S1.** Recombination events detected by concatenating the RNA-dependent RNA-polymerase sequences and the coat protein sequences.

**Fig. S2.** (A) Structure and quality prediction of totiviruses orthologs from the host. (B) structure comparison of Beauveria bassiana RNA virus 1 (totivirus) with host protein.

**Fig. S3.** Phylogenetic tree (species tree) produced by concatenating the RNA-dependant RNA-polymerase and the coat protein from partitiviruses and curvulaviruses used for aBSREL analysis. The curvulavirus sequences are highlighted in grey.

**Fig. S4.** Phylogenetic tree (species tree) produced by concatenating the RNA-dependant RNA-polymerase and the coat protein from chrysoviruses and curvulaviruses used for aBSREL analysis. The curvulavirus sequences are highlighted in grey.

**Fig. S5.** Phylogenetic tree produced by concatenating 70 proteins from the host and curvulavirus orthologs from the host.

**Table S1.** List of viruses used for PSI-BLAST search to retrieve RNA-dependent RNA-polymerase and coat proteins, and fungal proteins and the number of hits yielded using each of the sequences.

**Table S2.** List of viruses and their RNA-dependent RNA-polymerase and coat proteins accession numbers used in this study.

**Table S3.** Recombination events detected using the concatenated RNA dependent RNA polymerases and coat proteins.

**Table S4.** List of fungus proteins used in this study and their accession numbers.

**Table S5.** Nonsynonymous (*d_N_*) to synonymous (*d_S_*) substitutions ratio for the RNA dependent RNA polymerase (RdRps) and the coat proteins (CPs) from *Partitiviridae* and *Curvulaviridae*.

**Table S6.** Nonsynonymous (*d_N_*) to synonymous (*d_S_*) substitutions ratio for the RNA dependent RNA polymerase (RdRps) and the coat proteins (CPs) from *Chrysoviridae* and *Curvulaviridae*.

**Table S7.** List of the *Penicillium roqueforti* proteins used for BLASTp to retrieve homologous proteins within the fungi.

**Table S8.** List of fungi used to test transfer from virus to host.

**Table S9.** PSI-BLAST search results for homologs of *Penicillium roqueforti* proteins.

**Table S10.** Nonsynonymous (*d_N_*) to synonymous (*d_S_*) substitutions rates ratio (*ω*) calculation of the host proteins.

**Supplementary File 1.** Scripts written for the analyses and manipulations done in this study (HTML format).

## Acknowledgements

We thank Juan C. Muñoz-Sánchez, María J. Olmo-Uceda and João M.F. Silva for excellent suggestions and discussion. This work was supported by grant PID2019-103998GB-I00 funded by MCIN/AEI/10.13039/501100011033 and by Generalitat Valenciana grant CIPROM/2022/59 to S.F.E.

