## Supplementary Figures for "Evidence for gene transfer between mycoviruses and their host: *Curvulaviridae* as a case study"

### Fig S1

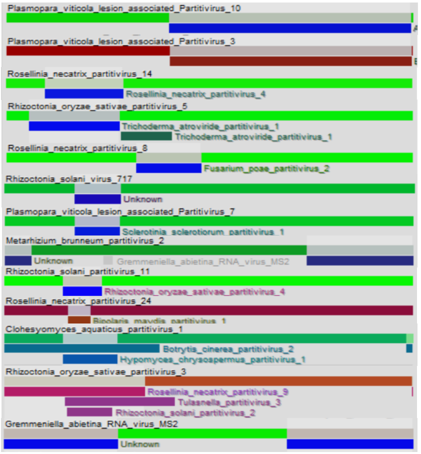

Plasmopara_viticola_lesion_associated_Partitivirus_10
Plasmopara_viticola_lesion_associated_Partitivirus_3
Rosellinia_necatrix_partitivirus_14
Rosellinia_necatrix_partitivirus_4
Rhizoctonia_oryzae_sativae_partitivirus_5
Trichoderma_atroviride_partitivirus_1
Trichoderma_atroviride_partitivirus_1
Rosellinia_necatrix_partitivirus_8
Fusarium_poae_partitivirus_2
Rhizoctonia_solani_virus_717
Unknown
Plasmopara_viticola_lesion_associated_Partitivirus_7
Sclerotinia_sclerotiorum_partitivirus_1
Metarhizium_brunneum_partitivirus_2
Unknown
Gremmeniella_abietina_RNA_virus_MS2
Rhizoctonia_solani_partitivirus_11
Rhizoctonia_oryzae_sativae_partitivirus_4
Rosellinia_necatrix_partitivirus_24
Bipolaris_maydis_partitivirus_1
Clohesyomyces_aquaticus_partitivirus_1
Botrytis_cinerea_partitivirus_2
Hypomyces_chrysospermus_partitivirus_1
Rhizoctonia_oryzae_sativae_partitivirus_3
Rosellinia_necatrix_partitivirus_9
Tulasnella_partitivirus_3
Rhizoctonia_solani_partitivirus_2
Gremmeniella_abietina_RNA_virus_MS2
Unknown

### Fig S2

**A**

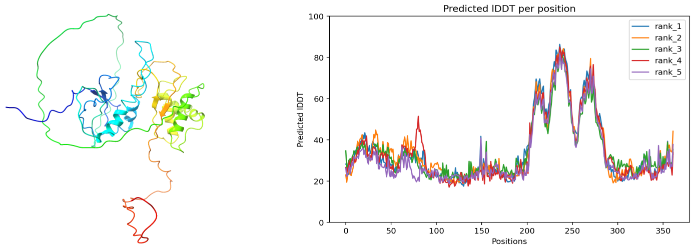


*Fusarium phyllophilum, hypothetical proteins (KAF5570093)*

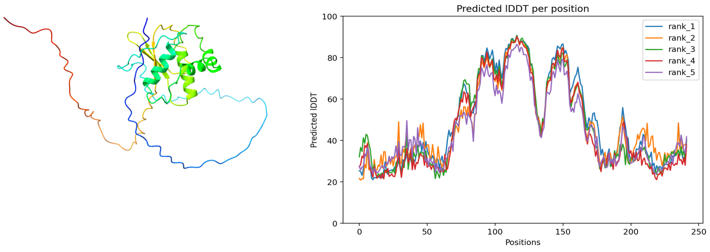


*Fusarium xylarioides, hypothetical protein (KAG5806886)*

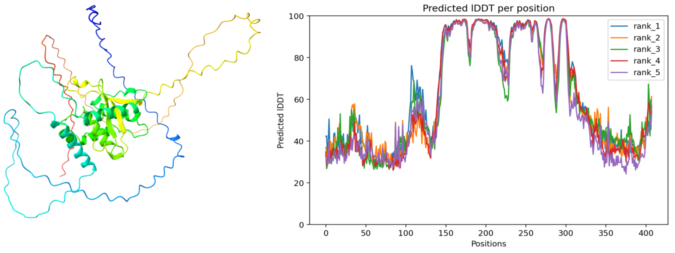


*Physcia stellaris, hypothetical protein (KAG6988846)*

**B**

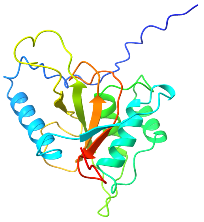

*Podospora anserina, UP,*
*(XP_001906714)*

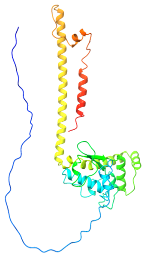

*Beauveria bassiana*
*RNA virus 1,*
*CP (AKC57300)*

(

### Fig S3

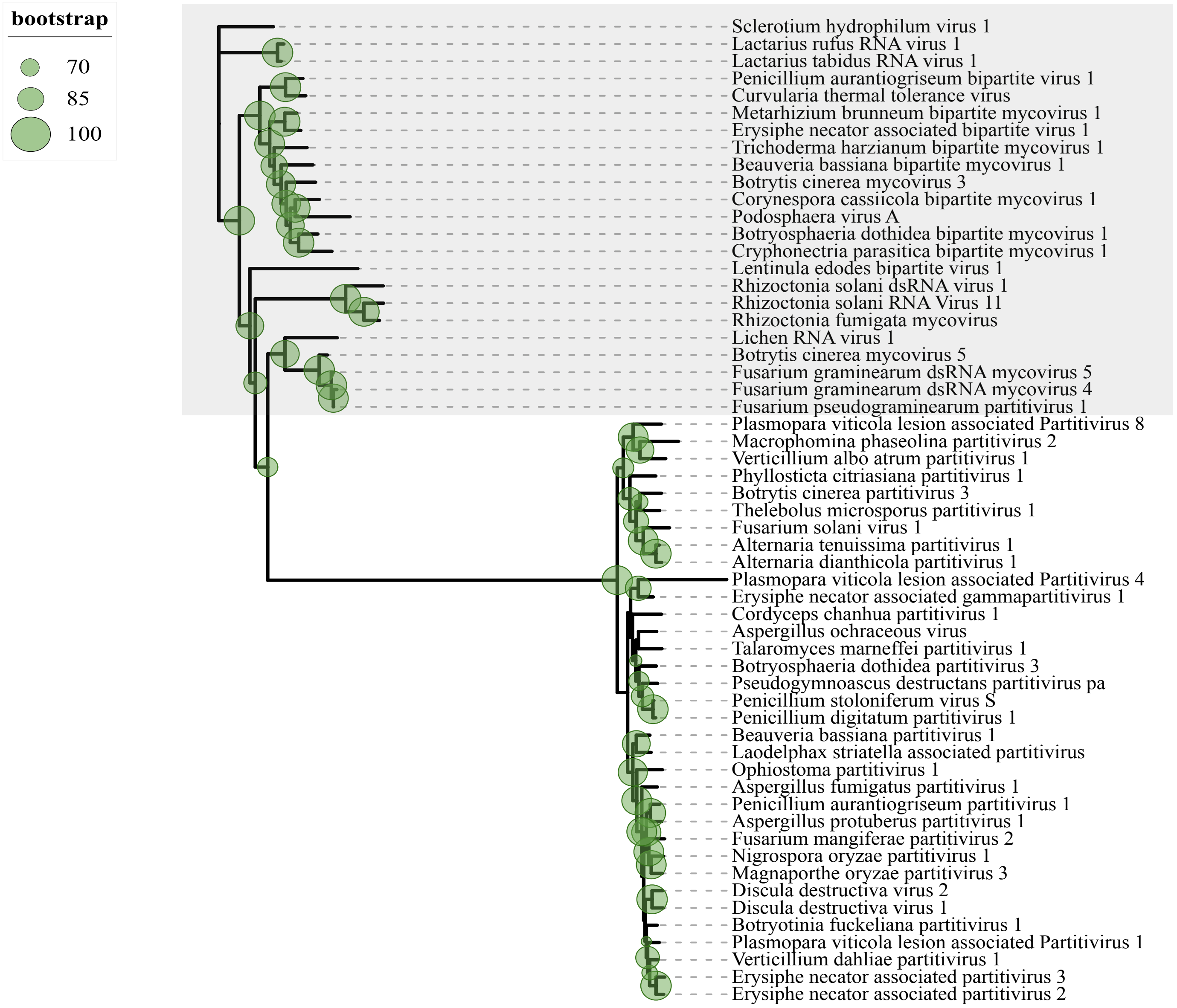
bootstrap
70
85
100
Sclerotium hydrophilum virus 1
Lactarius rufus RNA virus 1
Lactarius tabidus RNA virus 1
Penicillium aurantiogriseum bipartite virus 1
Curvularia thermal tolerance virus
Metarhizium brunneum bipartite mycovirus 1
Erysiphe necator associated bipartite virus 1
Trichoderma harzianum bipartite mycovirus 1
Beauveria bassiana bipartite mycovirus 1
Botrytis cinerea mycovirus 3
Corynespora cassiicola bipartite mycovirus 1
Podosphaera virus A
Botryosphaeria dothidea bipartite mycovirus 1
Cryphonectria parasitica bipartite mycovirus 1
Lentinula edodes bipartite virus 1
Rhizoctonia solani dsRNA virus 1
Rhizoctonia solani RNA Virus 11
Rhizoctonia fumigata mycovirus
Lichen RNA virus 1
Botrytis cinerea mycovirus 5
Fusarium graminearum dsRNA mycovirus 5
Fusarium graminearum dsRNA mycovirus 4
Fusarium pseudograminearum partitivirus 1
Plasmopara viticola lesion associated Partitivirus 8
Macrophomina phaseolina partitivirus 2
Verticillium albo atrum partitivirus 1
Phyllosticta citriasiana partitivirus 1
Botrytis cinerea partitivirus 3
Thelebolus microsporus partitivirus 1
Fusarium solani virus 1
Alternaria tenuissima partitivirus 1
Alternaria dianthicola partitivirus 1
Plasmopara viticola lesion associated Partitivirus 4
Erysiphe necator associated gammapartitivirus 1
Cordyceps chanhua partitivirus 1
Aspergillus ochraceous virus
Talaromyces marneffei partitivirus 1
Botryosphaeria dothidea partitivirus 3
Pseudogymnoascus destructans partitivirus pa
Penicillium stoloniferum virus S
Penicillium digitatum partitivirus 1
Beauveria bassiana partitivirus 1
Laodelphax striatella associated partitivirus
Ophiostoma partitivirus 1
Aspergillus fumigatus partitivirus 1
Penicillium aurantiogriseum partitivirus 1
Aspergillus protuberus partitivirus 1
Fusarium mangiferae partitivirus 2
Nigrospora oryzae partitivirus 1
Magnaporthe oryzae partitivirus 3
Discula destructiva virus 2
Discula destructiva virus 1
Botryotinia fuckeliana partitivirus 1
Plasmopara viticola lesion associated Partitivirus 1
Verticillium dahliae partitivirus 1
Erysiphe necator associated partitivirus 3
Erysiphe necator associated partitivirus 2

### Fig S4

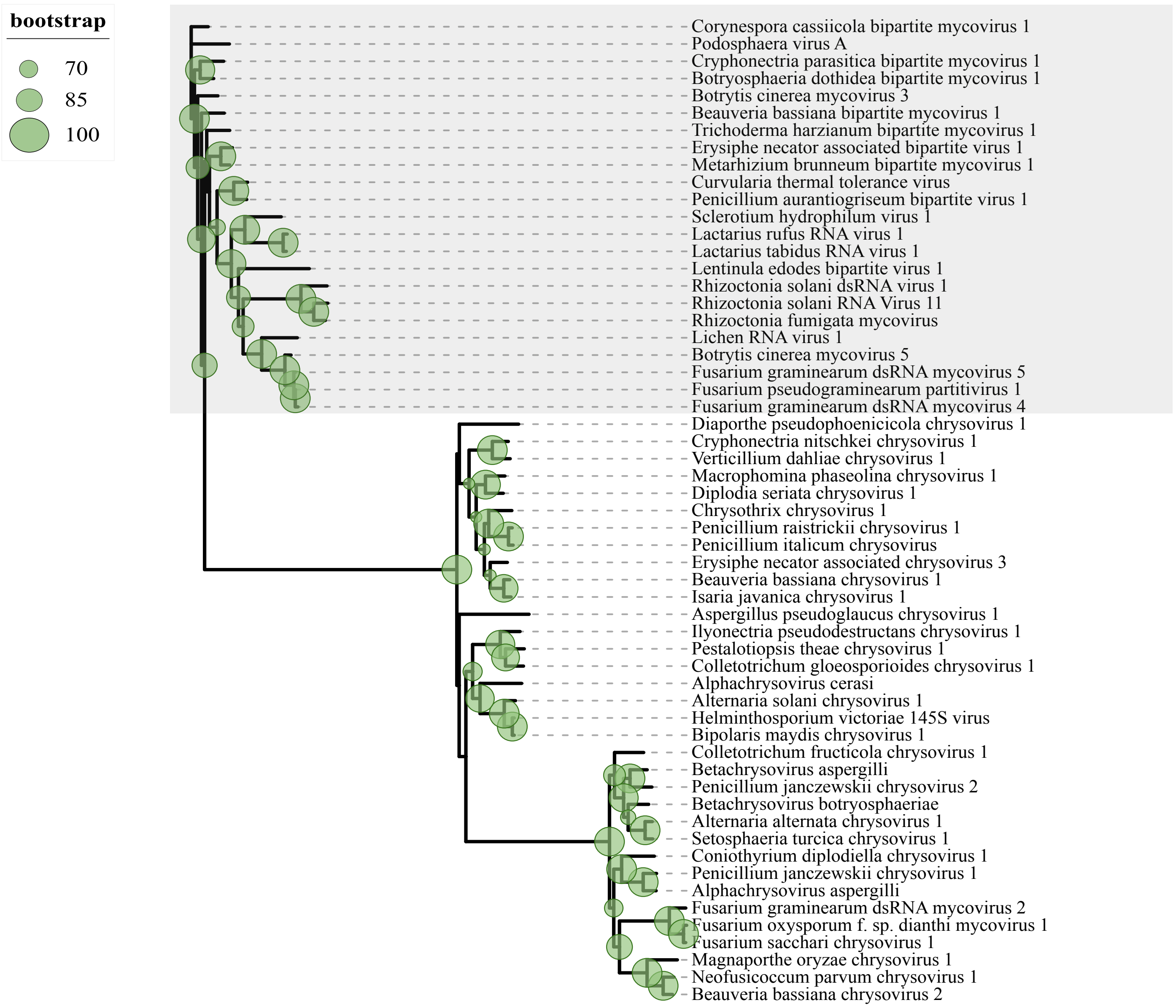
bootstrap
70
85
100
Corynespora cassiicola bipartite mycovirus 1
Podosphaera virus A
Cryphonectria parasitica bipartite mycovirus 1
Botryosphaeria dothidea bipartite mycovirus 1
Botrytis cinerea mycovirus 3
Beauveria bassiana bipartite mycovirus 1
Trichoderma harzianum bipartite mycovirus 1
Erysiphe necator associated bipartite virus 1
Metarhizium brunneum bipartite mycovirus 1
Curvularia thermal tolerance virus
Penicillium aurantiogriseum bipartite virus 1
Sclerotium hydrophilum virus 1
Lactarius rufus RNA virus 1
Lactarius tabidus RNA virus 1
Lentinula edodes bipartite virus 1
Rhizoctonia solani dsRNA virus 1
Rhizoctonia solani RNA Virus 11
Rhizoctonia fumigata mycovirus
Lichen RNA virus 1
Botrytis cinerea mycovirus 5
Fusarium graminearum dsRNA mycovirus 5
Fusarium pseudograminearum partitivirus 1
Fusarium graminearum dsRNA mycovirus 4
Diaporthe pseudophoenicicola chrysovirus 1
Cryphonectria nitschkei chrysovirus 1
Verticillium dahliae chrysovirus 1
Macrophomina phaseolina chrysovirus 1
Diplodia seriata chrysovirus 1
Chrysothrix chrysovirus 1
Penicillium raistrickii chrysovirus 1
Penicillium italicum chrysovirus
Erysiphe necator associated chrysovirus 3
Beauveria bassiana chrysovirus 1
Isaria javanica chrysovirus 1
Aspergillus pseudoglaucus chrysovirus 1
Ilyonectria pseudodestructans chrysovirus 1
Pestalotiopsis theae chrysovirus 1
Colletotrichum gloeosporioides chrysovirus 1
Alphachrysovirus cerasi
Alternaria solani chrysovirus 1
Helminthosporium victoriae 145S virus
Bipolaris maydis chrysovirus 1
Colletotrichum fructicola chrysovirus 1
Betachrysovirus aspergilli
Penicillium janczewskii chrysovirus 2
Betachrysovirus botryosphaeriae
Alternaria alternata chrysovirus 1
Setosphaeria turcica chrysovirus 1
Coniothyrium diplodiella chrysovirus 1
Penicillium janczewskii chrysovirus 1
Alphachrysovirus aspergilli
Fusarium graminearum dsRNA mycovirus 2
Fusarium oxysporum f. sp. dianthi mycovirus 1
Fusarium sacchari chrysovirus 1
Magnaporthe oryzae chrysovirus 1
Neofusicoccum parvum chrysovirus 1
Beauveria bassiana chrysovirus 2
